# Metabarcoding unsorted kick-samples facilitates macroinvertebrate-based biomonitoring with increased taxonomic resolution, while outperforming environmental DNA

**DOI:** 10.1101/792333

**Authors:** Lyndall Pereira-da-Conceicoa, Vasco Elbrecht, Andie Hall, Andrew Briscoe, Helen Barber-James, Benjamin Price

## Abstract

Many studies have highlighted the potential of DNA-based methods for the biomonitoring of freshwater macroinvertebrates, however only a few studies have investigated homogenisation of bulk samples that include debris to reduce sample-processing time. In order to explore the use of DNA-based methods in water quality assessment in South Africa, this study compares morphological and molecular-based identification of freshwater macroinvertebrates at the mixed higher taxon and mOTU level while investigating abundance and comparing mOTU recovery with historical species records. From seven sites across three rivers in South Africa, we collected a biomonitoring sample, an intensive-search comprehensive sample and an eDNA sample per site. The biomonitoring sample was picked and scored according to standard protocols and the leftover debris and comprehensive samples were homogenised including all debris. DNA-based methods recovered higher diversity than morphology, but did not always recover the same taxa, even at the family level. Regardless of the differences in taxon scores, most DNA-based methods except some eDNA samples, returned the same water quality assessment category as the standard morphology-based assessment. Homogenised comprehensive samples recovered more freshwater invertebrate diversity than all other methods. The eDNA samples recovered 2 to 10 times more mOTUs than any other method, however 90% of reads were non-target and as a result eDNA recovered the lowest target diversity. However, eDNA did find some target taxa that the other methods failed to detect. This study shows that unsorted samples recover the same water quality scores as a morphology-based assessment and much higher diversity scores than both picked and eDNA samples. As a result, there is potential to integrate DNA-based approaches into existing metrics quickly while providing much more information for the development of more refined metrics at the species or mOTU level with distributional data which can be used for conservation and biodiversity management.

## Introduction

The development of DNA-based identification methods for freshwater macroinvertebrates and their incorporation into biological monitoring programs is rapidly advancing, particularly due to the potential reductions in processing time, greater taxonomic resolution, and reduction in errors compared with current morphological monitoring methods (Hering et al., 2018). While morphological methods often have limited taxonomic resolution due to cryptic species, sex and life-stages, DNA-based methods can overcome these issues (Ekrem, Stur, & Hebert, 2010; Hebert, Ratnasingham, & de Waard, 2003; Park, Foottit, Maw, & Hebert, 2011; Venter & Bezuidenhout, 2016) when paired with suitable DNA reference libraries. For bulk-collected samples, sample sorting and subsequent morphological identification is time-consuming, and depending on the taxonomic expertise and available identification resources, the same samples can produce different results (Dickens & Graham, 2002; Haase et al., 2006) and up to a third of specimens and a fifth of taxa can be missed during the sorting stage (Haase, Pauls, Schindehütte, & Sundermann, 2010).

Within the last decade, a growing number of case-studies have highlighted the potential applications of DNA-based methods for the bioassessment of freshwater macroinvertebrates (Baird & Hajibabaei, 2012; Blackman et al., 2017; Carew, Pettigrove, Metzeling, & Hoffmann, 2013; Elbrecht & Leese, 2017; Elbrecht, Vamos, Meissner, Aroviita, & Leese, 2017; Elbrecht, Vamos, Steinke, & Leese, 2017a; Hajibabaei, Shokralla, Zhou, Singer, & Baird, 2011; Packer, Gibbs, Sheffield, & Hanner, 2009; Pauls et al., 2014; Yu et al., 2012), the majority of which pick invertebrates from the total sample as the first step. Only a few freshwater macroinvertebrate studies have investigated homogenisation, either of cleaned bulk samples where debris is removed (e.g. Andújar et al., 2018; Dowle, Pochon, Banks, Shearer, & Wood, 2016; Gardham, Hose, Stephenson, & Chariton, 2014) or bulk samples which still include some debris (Majaneva, Diserud, Eagle, Hajibabaei & Ekrem, 2018) in order to remove the specimen sorting stage and reduce sample-processing time.

In parallel, the use of environmental DNA (eDNA) is being explored to detect macroinvertebrates in freshwater environments. However, as eDNA is free of the organism and can be transported in flowing waters (Deiner & Altermatt, 2014), these data are spatially complex (Deiner, Fronhofer, Mächler, Walser & Altermatt, 2016). While eDNA-based studies have been shown to successfully detect macroinvertebrate community richness (Deiner et al., 2017; Deiner, Walser, Mächler & Altermatt, 2016; Li et al., 2018; Mächler, Little, Wüthrich, Alther & Fronhofer, 2019) some studies have shown a much lower detection level of invertebrates when compared to morphological methods (Hajibabaei et al., 2019; Macher et al., 2018). These differences can be a result of several factors including DNA shedding rates, persistence and movement in the environment (Barnes & Turner, 2016), sampling, laboratory and bioinformatic biases as outlined in Blackman et al (2019).

In addition to the sample processing and bioinformatic biases, DNA based methods are often limited by the lack of DNA reference libraries (Carew et al., 2017; Porter & Hajibabaei, 2018; Weigand et al., 2019), particularly in regions with high biodiversity and endemism (e.g. South Africa: Venter & Bezuidenhout, 2016). While large proportions of global biodiversity remain unknown (Stork, 2018), freshwater ecosystems are considered to be among the ecosystems most threatened by global climate change (Bates et al., 2008) and severe pressure from other anthropogenic impacts (Dallas & Rivers-Moore, 2014). Freshwater resources in Africa are highly impacted, and in South Africa, 57% of river and 75% of wetland ecosystems are highly threatened (Dallas & Rivers-Moore, 2014; Darwall, Smith, Tweddle, & Skelton, 2009; Driver, Sink, Nel, Holness, Van Niekerk, Daniels, Jonas, Majiedt, Harris, and Maze, 2012; Nel et al., 2011). A more in-depth knowledge of the current biodiversity would greatly improve our ability to inform decisions (Hamer, 2013) and is a critical factor hindering freshwater conservation in Africa (Barber-James & Pereira-da-Conceicoa, 2016).

While biological monitoring using freshwater macroinvertebrates is used worldwide, the indices are used at various taxon levels, with many indices only using family (or higher) level identification (e.g. South Africa: Dickens & Graham, 2002; Tanzania: Kaaya, Day, & Dallas, 2015; Namibia: Palmer & Taylor, 2004). However, such coarse taxonomic resolution has been shown to overlook the varying tolerances at generic or species levels, which can be used to make more well-informed water management decisions (e.g. Barber-James & Pereira-da-Conceicoa, 2016; Macher et al., 2016).

Within South Africa, rivers are currently monitored using the South African Scoring System (SASS version 5) protocol (Dickens & Graham, 2002) which uses freshwater macroinvertebrates as a measure of stream ecosystem health. Organisms are identified to a mixed taxon (typically family) level (Fig. S1) and are assigned a quality score based on pollution sensitivity (Dickens & Graham, 2002). The abundance of organisms is roughly estimated into categories (where 1 = 1 individual, A = 2 - 10, B = 10 - 100, C = 100 - 1000, D > 1000) and recorded on the scoring sheet. Abundance is not used in the SASS calculations, but is used in other indices, such as the Macroinvertebrate Response Assessment Index (MIRAI, Thirion, 2007). SASS returns three principal indices: SASS Score (the total of quality scores for all taxa found in a sample), Number of Taxa and ASPT (Average Score Per Taxon = SASS Score divided by Number of Taxa) (Dickens & Graham, 2002). These scores are then standardised across South Africa using biological bands (A-E) calibrated for each ecoregion (Dallas, 2007; Dallas & Day, 2007; Kleynhans, Thirion, & Moolman, 2005; Omernik, 1987). Identification at the SASS level (i.e. the mixed higher taxon levels) is too coarse for biodiversity and ecological impact assessment, and should only be used as a red-flag indicator of changes in water quality of a monitored site (Dickens & Graham, 2002), however due to the lack of alternative methods SASS is frequently misused in southern Africa (Barber-James & Pereira-da-Conceicoa, 2016). Data should be interpreted in relation to season of collection as some natural variation will occur throughout the year (Dickens & Graham, 2002) and seasonal data is required to capture a holistic view of the taxa occurring at a site.

In order to explore the use of DNA-based methods of assessment in South Africa, this study compares morphological and molecular-based identification of freshwater macroinvertebrates at the mixed higher taxon SASS level and at the level of the molecular operational taxonomic unit or “mOTU”. DNA metabarcoding of picked SASS samples, leftover SASS debris (after picking), intensive-search comprehensive samples and filtered water samples (eDNA) were compared with morphology. Three main aims were investigated: 1) which of the sampling and processing methods (picked SASS, leftover SASS, comprehensive sample or eDNA), recover the most similar SASS level taxa and scores as morphology; 2) The correlation between the abundance of picked samples and relative abundance of DNA sequence reads; and 3) Does DNA based mOTU recovery reflect historical records of species known from the region.

## Materials and Methods

### Study sites and sampling strategy

Samples were collected from seven sites across three Eastern Cape rivers, three sites along each of the Elandsbos (“E”) and Tyume tributary (“H”) rivers and one site on the Berg (“CD”) River (Fig. S2). River sites occurred across three ecoregions (Ecological Level 1 and geomorphological zone, (Dallas, 2007; Dallas, 2005) namely: Southeastern Coastal Belt (E = Elandsbos), upper Southeastern Uplands (H = Tyume tributary) and Southern Folded (CD = Berg). All sites sampled were selected based on the expected condition of good quality water in a natural or unmodified river condition.

At each site environmental DNA (eDNA) was collected from surface water using a sterile 1L bucket from five locations within a 5m radius of the sample site, and mixed in a sterile 20L bucket. Water was then filtered through three 0.22 μm polyethersulfone filters (Sterivex-GP) using a 50ml syringe. At each site water was filtered until the filter blocked or 2L was reached (total volume filtered ranged between 0.6 - 2L). Each filter was preserved onsite with 96% ethanol and kept refrigerated where possible.

Following eDNA sampling a SASS sample was taken by an accredited river health practitioner, following the SASS protocol using a 30×30cm framed standard kicknet with a 1mm mesh: a 2 minute kick-sample of stones-in-current across the river, 1 minute kick-sample of stones-out-of-current, 2m total marginal vegetation sweep and a 1 minute stir and sweep of gravel, sand and mud biotope (Dickens & Graham, 2002). SASS samples, including all debris (i.e. the entire kicknet contents), were preserved in 96% ethanol while in the field. As SASS samples were identified in the laboratory, rather than live in the field, the results are not comparable with other SASS data from these sites. In order to provide a more comprehensive assessment of the biodiversity at each site, whole community samples (here termed “Comprehensive”) were collected for 30 minutes using a 30×30cm framed kicknet with a 250um mesh. The whole sample, including all debris was preserved in 96% ethanol while in the field.

### Sample processing (Fig. 1)

SASS samples were scored by a single individual and identified according to SASS protocol (i.e. a total time of 45 minutes per sample for identification to family level). Invertebrates from each sample were picked, morpho-sorted after scoring and then identified further if possible and counted (here termed “SASS picked”). A representative individual of each morpho-species from the SASS picked samples was removed as a voucher specimen. Vouchers were imaged before being placed in ATL buffer and proK overnight at 56 °C and DNA was extracted using the Qiagen BioSprint 96 DNA Blood Kit or the DNeasy Blood & Tissue Kit.

**Fig. 1.**
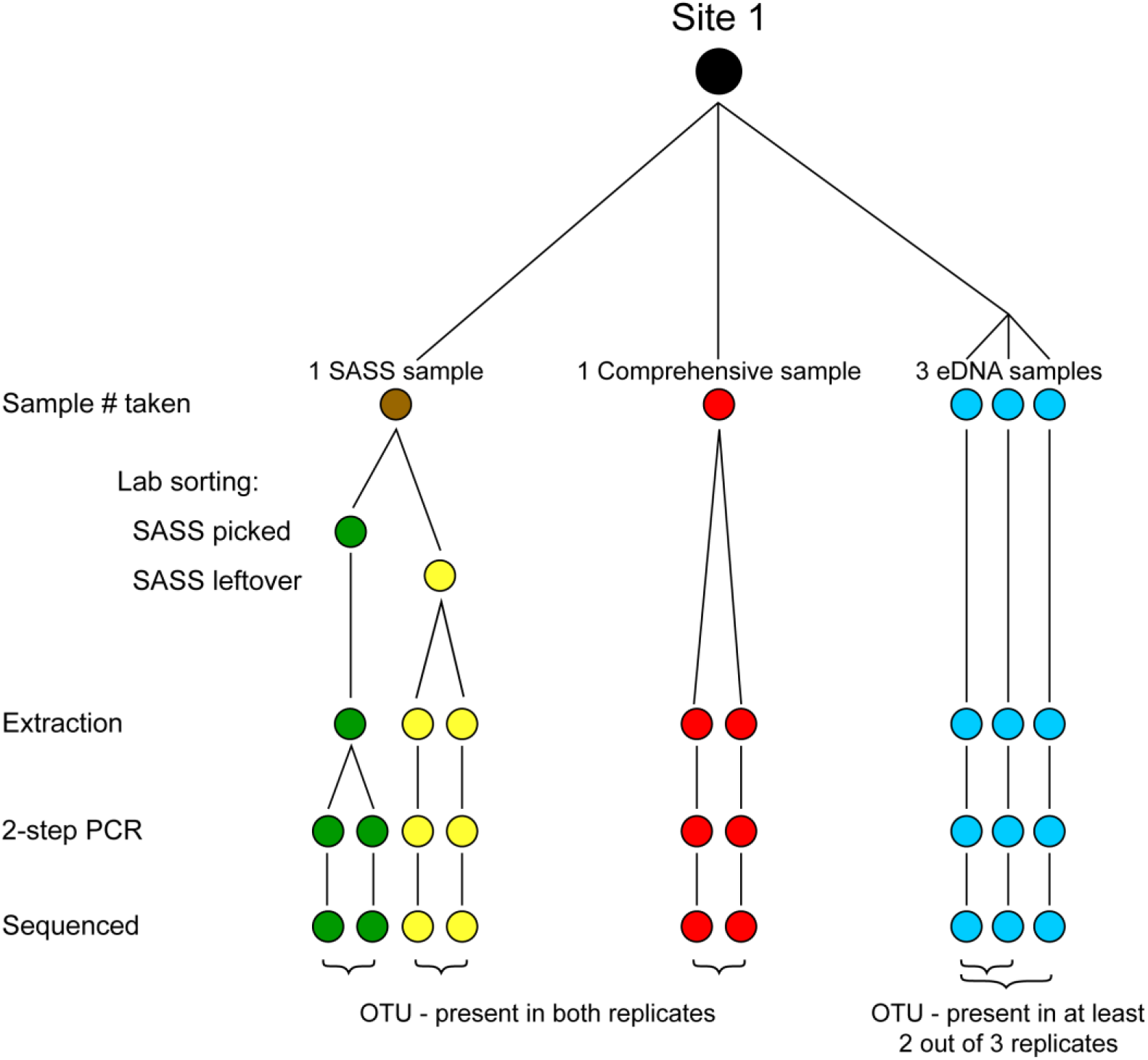
Sampling strategy per river site showing laboratory procedures and the corresponding technical replicates used for the study.

The remainder of the picked morpho-species (SASS picked) samples were placed in 2ml tubes and dried using a DNA SpeedVac dna120 at high heat (65 °C) for one large sample (H4) and medium heat (43 °C) for all other samples. The SASS picked samples were then homogenised with three glass beads per 2 ml tube using the Qiagen TissueLyser II (30 Hz for 1 minute) and digested overnight in 10:1 solution of ATL buffer and proK. Following digestion, three 200ul sub-samples were taken and extracted using the DNeasy Blood & Tissue kit with a double final elution resulting in 300ul DNA for each sub-sample. These sub-samples were then combined into a single extract of 900ul for each SASS picked sample from each site.

The remainder of the SASS samples, consisting of debris and some macroinvertebrates (here termed “SASS leftover”) and the Comprehensive samples were homogenised in ethanol for up to 10 seconds using a consumer blender (Breville VBL062, 300 W, 600 ml). The blender blades and container were sterilized using 12% industrial bleach between samples. Both the blended Comprehensive and SASS leftover samples were then sub-sampled with two replicates of 10g homogenate per sub-sample. Samples were dried overnight at room temperature and DNA was extracted using concentrated proK (20mg/ml) with overnight digestion following Ransome et al. (2017) and then the Qiagen DNeasy PowerMax Soil kit with a final DNA elution of 5ml per sub-sample. For the eDNA samples, the Sterivex filters (three replicates per site, Fig. 1) were dried and DNA extracted using the Qiagen DNeasy PowerWater Sterivex kit, following the alternative method (using a microcentrifuge and not a vacuum manifold), and excluding the 90°C incubation and powerbead steps. All DNA concentrations were quantified using a NanoDrop 8000 (Thermo Fisher Scientific) and the concentration of DNA was adjusted to 25 ng/μl in the downstream PCRs.

### Voucher barcoding

Voucher barcoding followed standard in-house protocols using LCO1490 and HCO2198 primers (Folmer, Black, Hoeh, Lutz, & Vrijenhoek, 1994). Each reaction consisted of 1mM total dNTPs, 3mM MgCl_2_, 1.25u Bio-Taq DNA polymerase (Bioline), 0.1μM each primer and 1x reaction buffer. Cycling conditions were: initial denaturation 94°C for 1min followed by 35 cycles of 94°C for 30s, 50°C for 30s and 72°C for 30s, with a final elongation of 10min at 72°C. PCR products were visualised using gel electrophoresis, purified using Agencourt AMPure XP beads and then sequenced bi-directionally using BigDye terminator reaction mix v3.1 in a 3730xl DNA analyser (Applied Biosystems) at the NHM sequencing facility. Voucher data and corresponding sequences were uploaded to the Barcode of Life Data System (BOLD) website (Ratnasingham & Herbert, 2007) and can be found under the project name “BEISA” in the public data portal.

### DNA metabarcoding

DNA metabarcoding followed the 2-step PCR approach outlined in Elbrecht and Steinke (2018) using the BF2 / BR2 freshwater macroinvertebrate fusion primer sets developed for the Cytochrome c oxidase subunit I (COI) gene (Elbrecht & Leese, 2017). Each SASS picked extraction was amplified in duplicate, for other methods each technical replicate was independently amplified (two for Comprehensive and SASS leftover, and three for eDNA samples, Fig. 1).

Two-step PCRs were performed on each sample replicate using the Qiagen Multiplex PCR Plus Kit with 0.5 uM of each primer in a final volume of 25ul. PCRs were run on Techne™ Prime Elite Thermal Cyclers (Thermo Fisher Scientific) with the following conditions, PCR 1: 94 °C for 5 min; 25 cycles of 94 °C for 30s; 50 °C for 30s; 65 °C for 50s; and final extension at 65 °C for 5 min. The eDNA samples used 27 cycles rather than 25. Untailed BF2 / BR2 primers were used for PCR 1. Then 1ul of amplicon from the initial PCR was used as a template for PCR 2 under the following conditions: 94 °C for 5 min; 13 cycles of 94 °C for 30s; 50 °C for 30s; 65 °C for 2 min; and final extension at 65 °C for 5 min using the tailed BF2 / BR2 fusion primers (Elbrecht & Leese, 2017). One negative control was used in both PCR steps, PCR products were visualised using gel electrophoresis.

Following the second PCR, the tagged amplicons were purified using Agencourt AMPure XP beads at 0.8x ratio. The eDNA sample from site E3 was gel cut using the QIAquick Gel Extraction Kit to remove a non-target band. Following cleanup the DNA concentration of each individual library was measured with a SPECTROstar Nano (BMG Labtech) and then equimolar pooled with negative controls added at the maximum volume added for any single library (15ul).

The size-corrected concentration of the pooled libraries was determined following analysis with an Agilent 2200 Tapestation system and Qubit 2.0 Fluorometer (Invitrogen). The pool was loaded onto an Illumina MiSeq at 9pM, with 5% Phi-X, using a 600 cycle V3 kit with 300 bp paired end sequencing (index read steps skipped).

### Bioinformatics

Bioinformatics processing was performed using JAMP v0.66 (http://github.com/VascoElbrecht/JAMP) with detailed scripts being available as supporting information (Scripts S1). Reads were demultiplexed using JAMP, and paired-end reads were merged using Usearch v8.1.1861 (Edgar, 2013) with relaxed settings to maximise the amount of reads merged (allowing up to 99 mismatches in the overlapping region). Where necessary, the reverse complement of the reads was generated, to ensure all sequences are present in the same orientation. BF2 and BR2 primers were then removed using cutadapt 1.18 with default settings (Martin, 2011), discarding reads where the primer sequences remained undetected. Only sequences of 411 to 431 bp were used for further analysis (filtered with cutadapt). Low-quality sequences were then filtered from all samples, using fastq_filter with maxee = 1 (Edgar & Flyvbjerg, 2015). Sequences from all samples were then pooled, dereplicated (minuniquesize = 2), and clustered into molecular operational taxonomic units (mOTUs), using cluster_otus with a 97% identity threshold (Edgar, 2013) which includes chimera removal. Pre-filtered reads for all samples were de-replicated again, but singletons were included to maximise the information extracted from the sequence data. Sequences from each sample were matched against the mOTUs with a minimum match of 97% using usearch_global.

For each sample only mOTUs with a read abundance above 0.01% in at least two sequencing replicates were considered for downstream analyses using a 3% divergence threshold which is consistent with other studies and observations of real invertebrate communities (e.g., Elbrecht, Peinert, & Leese, 2016; Elbrecht, Vamos, Steinke, & Leese, 2017b; Hajibabaei, Janzen, Burns, Hallwachs, & Hebert, 2006).

Taxonomy was assigned to remaining mOTUs using an R script to search against both BOLD and NCBI databases. The taxonomy assigned mOTU table was then filtered by Phylum to include targeted taxa (Arthropoda, Annelida, Porifera, Coelenterata, Turbellaria, Hydracarina, Gastropoda and Pelecypoda), taxonomy was further validated and checked, removing any terrestrial invertebrates. Any conflicting assignments between BOLD and NCBI were handled individually and unresolved cases were removed. A rough guideline for taxonomy assignment using percent similarity was used, where a hit with 98% similarity was used for species, 95% for genus, 90% for family and 85% for order levels. UpSetR (Conway, Lex, & Gehlenborg, 2017; Lex et al., 2014) was used to visualise the number of shared mOTUs / families between each metabarcoding sampling method across all sites.

The number of SASS level family taxa detected by each DNA metabarcoding sampling method was then compared to morphology. Within the SASS picked samples singletons were artificially removed from the comparison as they were used for voucher sequencing (i.e. not available for metabarcoding in the SASS picked sample). The percentage of taxon overlap (at SASS level) of DNA methods with morphology was then calculated using R (version 3.5.3; R Core Team, 2019).

### Water Quality Assessment

For each site, the ecoregion classification was determined based on Kleynhans et al. (2005), and the total Taxon Score and Average Score Per Taxon (ASPT) values calculated for each method (morphology, SASS picked, SASS leftover, Comprehensive and eDNA) and plotted against the biological band for that region. The SASS picked and SASS leftover taxon lists for each sample were also combined and deduplicated to simulate a full SASS sample being processed as a single unit.

Although SASS does not rely on measures of abundance, this is recorded and can be used for other indices, such as the Macroinvertebrate Response Assessment Index (MIRAI, Thirion, 2007). At the family (or higher taxon) level, the relationship between SASS-level morphologically detected abundance and relative read abundance was explored using linear regression.

### Historical species records

The taxa previously recorded for each river / sampling area were extracted from the database of the Albany Museum, Makhanda (previously known as Grahamstown), which houses the largest freshwater invertebrate collection in Africa, including from the sites sampled in this study, and compared to the target mOTUs recovered by molecular methods. Only records identified at the genus or species level were used for comparison. Where available, data on the number of described species for the broader region (Eastern Cape / South Africa / southern Africa as applicable) was added to the analysis.

## Results

### Sequencing statistics

The MiSeq run yielded 16 million reads from the 68 tagged samples (raw data available from https://doi.org/10.5281/zenodo.3462633). After library demultiplexing, an average of 221,630 (SD = 69,340) read pairs were retained. After bioinformatic processing, a total of 17,660 mOTUs were detected which was then reduced to 5,117 mOTUs that were present in more than one technical replicate. Taxonomy was assigned using BOLD and NCBI, the mOTU table was filtered according to freshwater macroinvertebrate fauna listed in the SASS protocol resulting in a total of 404 “target” mOTUs being found across all samples. A regression comparing volume of water and number of target mOTUs found per eDNA Sterivex filter (N = 21) found a slightly negative but significant correlation, with a low adjusted R^2^ value (adj. R^2^ = 0.1854, slope = −0.00345, P = 0.02936).

The proportion of reads lost from each processing step to the 404 target taxa found across samples and sites are shown in Fig. 2. SASS picked samples represented the “cleanest” samples, with the lowest proportions of reads discarded (mostly under 20% discarded), Comprehensive and SASS leftovers showed similar results (ca. 40 - 50% discarded), while over 90% of reads were discarded for the eDNA samples (Fig. 2).

**Fig. 2.**
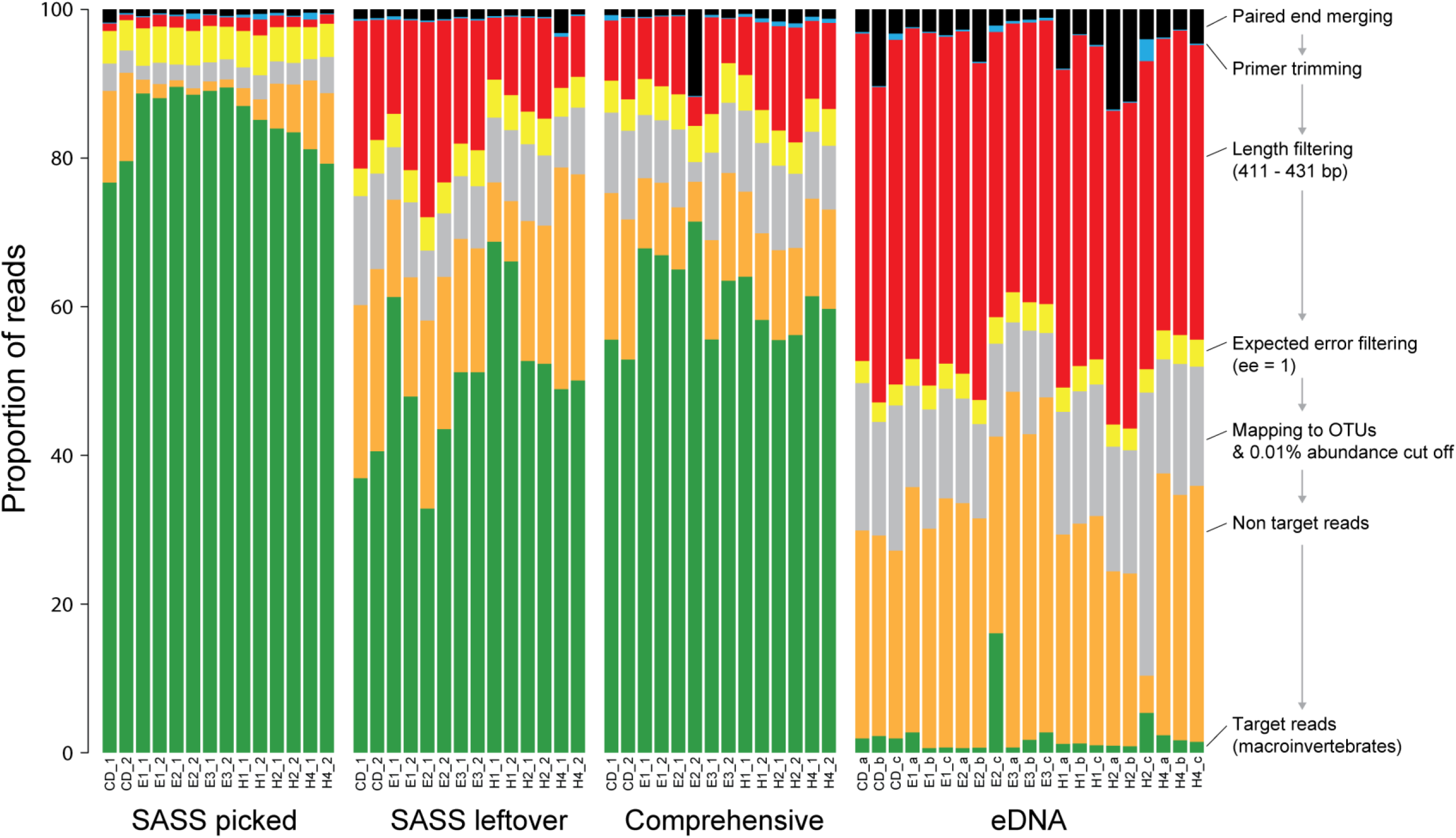
Proportion of reads lost during each processing step, an overview of sequences discarded from raw data, bioinformatics processing and non-target hits (e.g. bacteria) compared to the target macroinvertebrate taxa (green).

### Overlap between DNA methods

When considering the number of mOTUs shared between methods across all sites, only 57 of the 5,117 mOTUs (0.01%) found were shared between all four methods (Fig. 3a). As expected, SASS picked returned mostly arthropods (237 mOTUs), 76% of which were also present in the SASS leftover sample and 84% of SASS picked mOTUs were shared with the Comprehensive sample (Fig. 3a). The eDNA samples showed the least overlap with other methods, but returned the highest number of mOTUs (3,446) most of which were unique (84%). The Comprehensive (1,249 mOTUs) and SASS leftovers (1,780 mOTUs) shared 841 mOTUs (Fig. 3a).

**Fig. 3.**
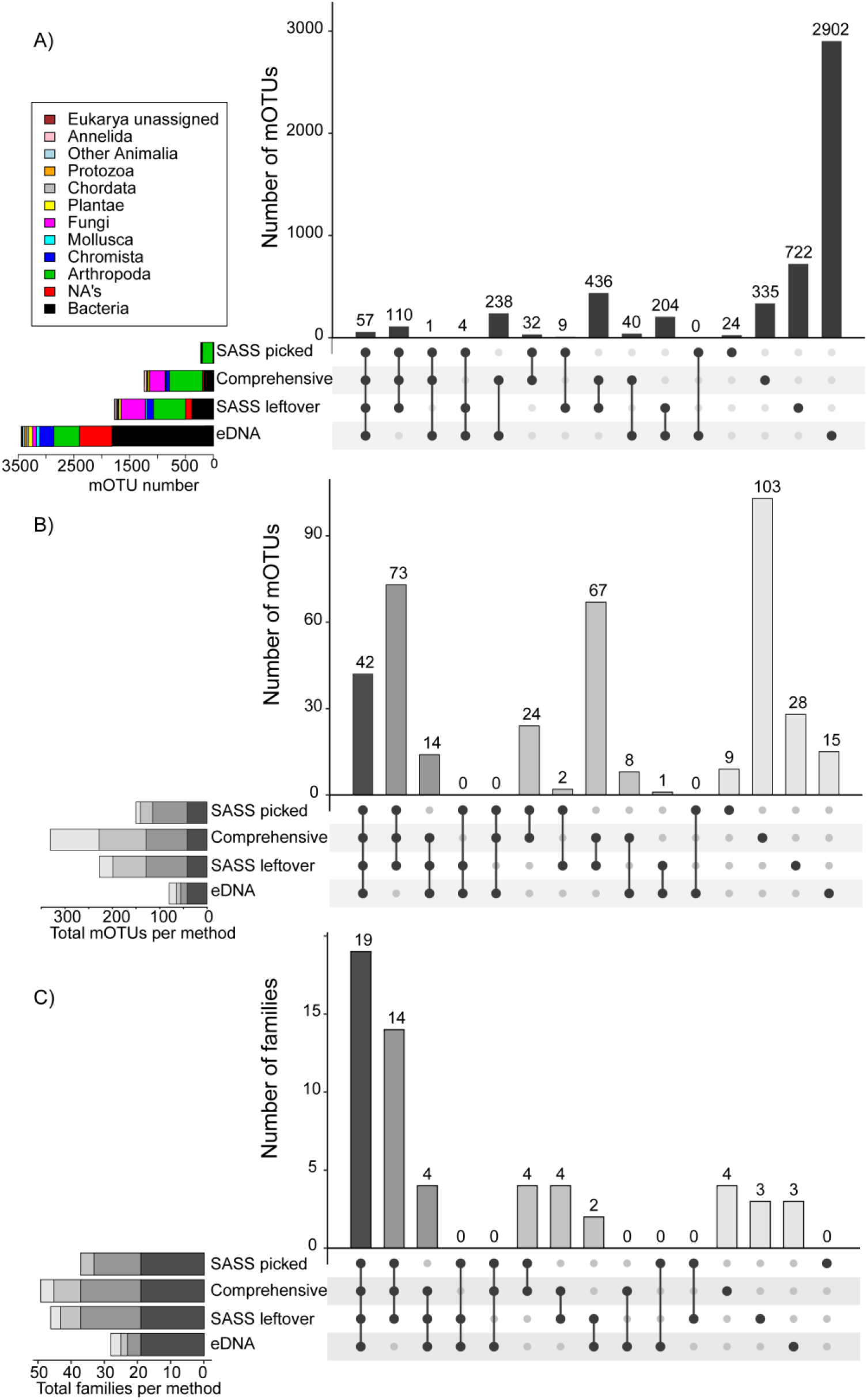
UpSet bar plot showing shared mOTUs between each sampling method across all sites. (A) For the 5117 subset mOTUs and (B) of the 404 mOTUs representing target freshwater invertebrates listed in the SASS protocol. C). UPset plot showing overlap of family level taxa between DNA based methods.

For targeted SASS mOTUs, only 42 of 404 mOTUs (0.1%) were shared between all four methods (Fig. 3b). SASS leftover (227 mOTUs) and Comprehensive samples (331 mOTUs) overlapped with SASS picked by 78% and 93% respectively, and shared 196 mOTUs. The eDNA samples recovered 80 target mOTUs, 19% of which were unique and not found by any other method. The Comprehensive sample had the highest number of unique mOTUs (103) corresponding to 26% of all target mOTUs found (Fig. 3b).

At the SASS mixed taxon (mostly family) level 19 out of 57 taxa were shared between all four methods (Fig. 3c). The three kick-net based methods (SASS picked, SASS leftover and Comprehensive) shared a further 14 taxa. The eDNA samples shared another 2 taxa with SASS leftovers, and each method, except SASS picked, found 2-3 unique taxa (Fig. 3c).

### Taxonomic identification at family level

At the SASS mixed taxon level, morphological based identification resulted in 51 taxa (Table S1) with an average of 21.86 ± 2.97 taxa at each site. SASS picked was most similar to morphology (mean = 20.86 ± 3.63), as expected given the morphotaxa came from these samples. Four SASS taxa were not recovered by DNA metabarcoding across the SASS picked samples: Ancylidae (Gastropoda) which was missing from 2/7 samples, Hydroptilidae (Trichoptera) missing from 2/7 samples, Gyrinidae and Chironomidae each missing from one sample. Seven families that were not recorded by morphology in some SASS picked samples were recovered by DNA metabarcoding: Caenidae, Notonemouridae, Philopotamidae, Pisuliidae, Teloganodidae and Tipulidae, potentially a result of sequencing gut contents of other invertebrates.

Comprehensive sample DNA metabarcoding generally found more target families than all the other methods (mean = 25.71 ± 4.75), and recovered nearly 4 more families than morphology (Fig. 4a) on average across sites. All other methods found fewer target families than morphology on average. SASS leftover recovered between 14 - 26 target families (mean = 19.14 ± 3.72) while taxon recovery from the eDNA samples was consistently lower than other methods, between 5 - 16 target families (mean = 10.29 ± 3.82) (Fig. 4a).

**Fig. 4.**
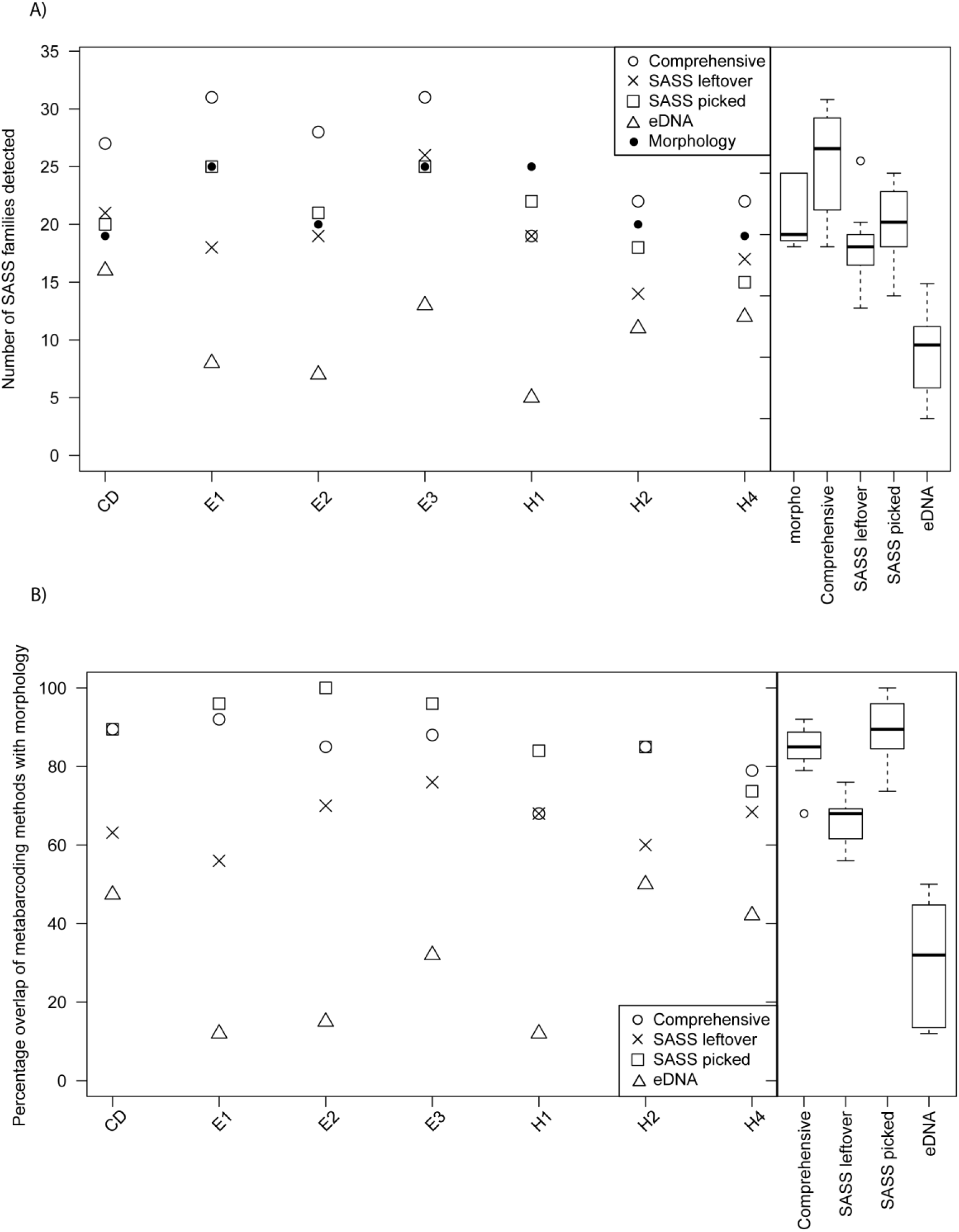
(A) number of SASS mixed taxon level taxa detected by morphological SASS and DNA-based methods (eDNA, SASS picked, SASS debris and Comprehensive debris) across sites. (B) percentage overlap of the taxa detected by DNA-based methods (eDNA, SASS picked, SASS debris and Comprehensive debris) with morphology.

Despite the Comprehensive method finding more family taxa than had been observed using morphology, the method did not recover all the same taxa with a 68 - 92 % overlap with morphologically identified taxa (failing to detect between 2 - 8 taxa, mean = 3.57 ± 2.07) (Fig. 4b). SASS picked samples showed a 74 - 100 % overlap with morphology, failing to detect between 0 - 5 taxa (mean = 2.29 ± 1.80) across all samples. SASS leftover failed to detect an average of 34% (6 - 11 taxa, mean = 7.43 ± 1.81), while eDNA failed to detect an average of 70% (10 - 22 taxa, mean = 15.57 ± 5.32) of morphologically identified taxa, respectively (Fig. 4b).

Water quality assessment metrics calculated for SASS-level morphology and DNA metabarcoding data were generally similar across methods when interpreted using Ecoregion Level 1 classified biological bands (Dallas, 2007), except for some eDNA samples (Fig. 5a-c). The total taxon scores varied considerably across sites sampled, even based on morphology, however ASPT was less variable and is recommended as the common SASS index of choice (Dickens & Graham, 2002), this is particularly evident in the Craigdoone site (Fig. 5a), where the total taxon scores differed by over 75 units while ASPT remained very similar between methods.

**Fig. 5.**
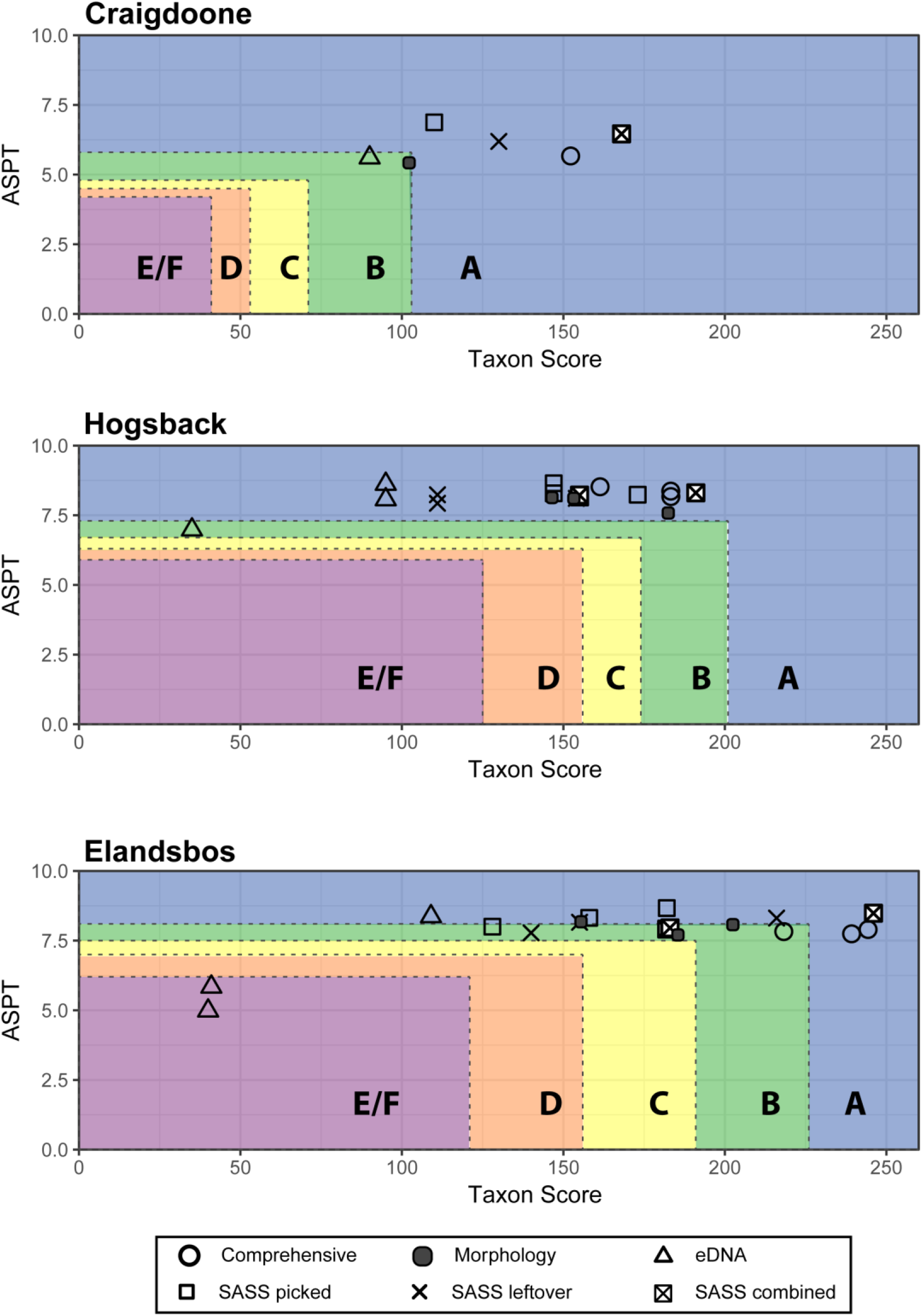
Plot of Taxon Score and ASPT scores, showing results for each site according to method. Biological bands are shown for each region, calculated through the intersection of total score and ASPT: A = natural (blue), B = good (green), C = fair (yellow), D = poor (red) and E/F = seriously modified (purple).

The relative proportion of reads assigned to each family was positively correlated with abundance when all sites were combined into a single regression (Fig. 6) with a significant but moderate adjusted R^2^ value (R^2^ = 0.557, p < 0.0001).

**Fig. 6.**
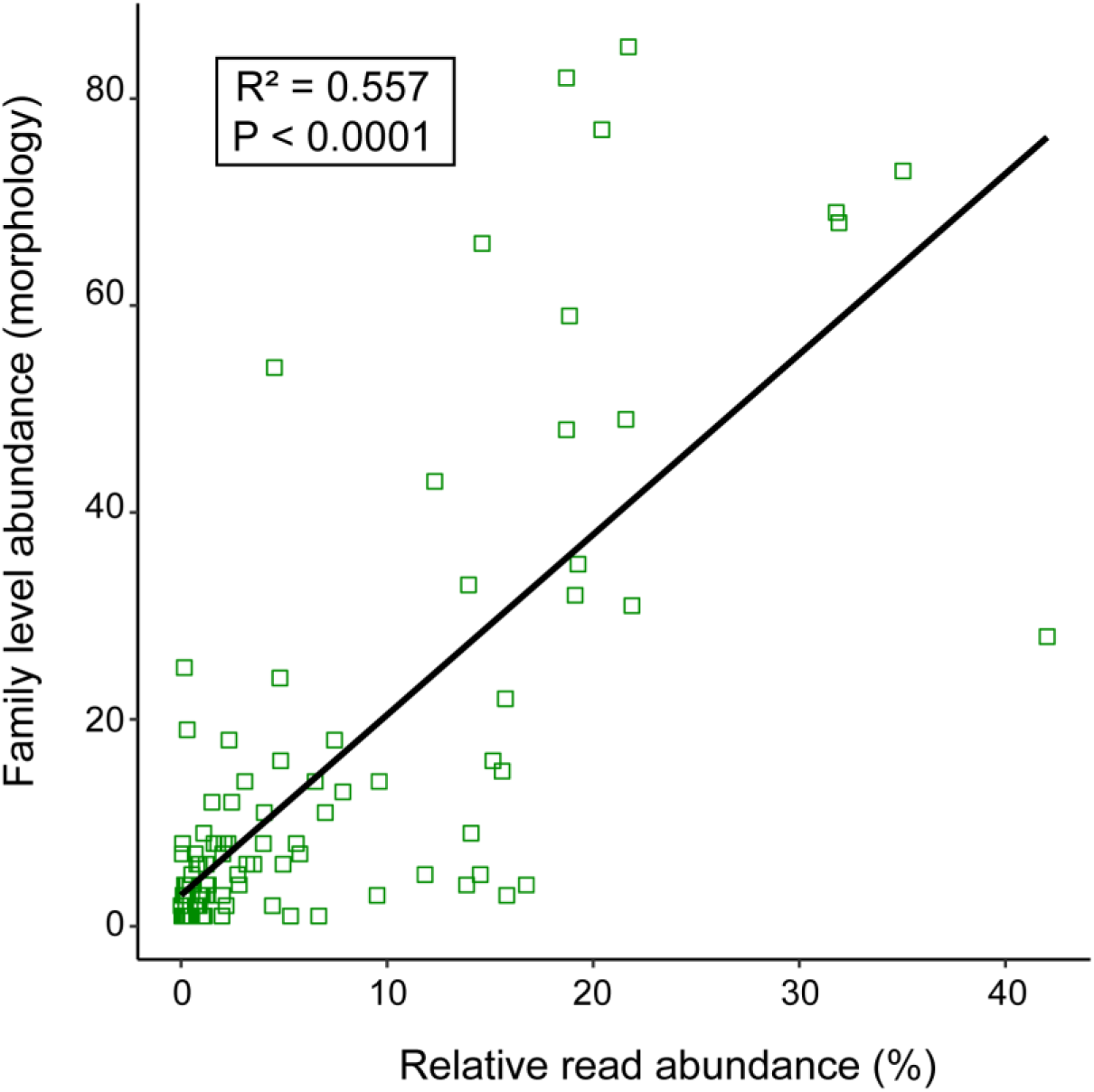
Relative abundance of sequences from SASS picked samples plotted against number of specimens morphologically identified at SASS-level for all sites sampled.

### Taxonomic identification at morphotaxon level

A total of 319 morphotaxa were picked from the SASS samples, identified to mixed taxon level and sanger sequenced to be used as vouchers. Of these, duplicate species as identified by COI sequences and non-target taxa (terrestrial invertebrates) were removed, leaving 254 unique freshwater macroinvertebrate morphotaxa. Voucher sequences were uploaded to BOLD before taxonomy assignment on the metabarcode data commenced. Morphotaxa were identified mostly to family and genus level using the Guide to the Freshwater Invertebrates of Southern Africa Series (Water Research Commission, specific references in Table 1), while NCBI and BOLD were used to confirm identifications and help infer some species names.

**Table 1.** Number of mOTUs recovered for each family with DNA-based methods, compared to Albany Museum historical specimen records of species found at the rivers sampled in this study. Regional records for families with number of mOTUs >5 and for those with regional species information readily available. Regional information available include Eastern Cape (EC) sometimes with Western Cape (WC), South Africa (SA) and southern Africa.

| **Taxon** | **# mOTUs** | **# Historical records** | **#species known from river sites** | **Regional** |
| --- | --- | --- | --- | --- |
| Chironomidae | 101 | 2 | 30 | EC<sup>1</sup> |
| Oligochaeta | 42 | 1 | 29 | EC<sup>2</sup> |
| Simuliidae | 34 | 12 | 23 | EC<sup>3</sup> |
| Baetidae | 24 | 17 | 37 | southern Africa<sup>4</sup> |
| Potamonautidae | 23 | 2 | 14 | SA<sup>5</sup> |
| Notonemouridae | 13 | 4 | 8 | EC<sup>6</sup> |
| Hydracarina | 10 | 8 | 23 | EC<sup>7</sup> |
| Platycnemididae | 10 | 1 | 2 | EC<sup>8</sup> |
| Coenagrionidae | 9 | 10 | 18 | EC<sup>8</sup> |
| Gyrinidae | 9 | 4 | 24 | EC<sup>9</sup> |
| Leptophlebiidae | 9 | 5 | 20 | southern Africa<sup>4</sup> |
| Leptoceridae | 8 | 6 | 36 | WC/EC<sup>10</sup> |
| Ceratopogonidae | 5 | 2 | 142 | southern Africa<sup>11</sup> |
| Dytiscidae | 5 | 3 | 27 | EC<sup>12</sup> |
| Elmidae | 5 | - | 5 | EC<sup>13</sup> |
| Scirtidae | 5 | 1 | ? | unknown<sup>14</sup> |
| Coelenterata | 4 | - |  |  |
| Corydalidae | 4 | 3 |  |  |
| Gomphidae | 4 | 5 | 5 | EC<sup>8</sup> |
| Hydropsychidae | 4 | 5 |  |  |
| Notonectidae | 4 | 1 |  |  |
| Sphaeriidae | 4 | 1 |  |  |
| Teloganodidae | 4 | 2 | 2 | EC<sup>4</sup> |
| Tipulidae | 4 | 2 |  |  |
| Amphipoda | 3 | - |  |  |
| Athericidae | 3 | 1 |  |  |
| Corixidae | 3 | 2 |  |  |
| Naucoridae | 3 | 1 |  |  |
| Pisuliidae | 3 | 1 |  |  |
| Porifera | 3 | - |  |  |
| Sericostomatidae | 3 | 2 |  |  |
| Turbellaria | 3 | - |  |  |
| Caenidae | 2 | 2 |  |  |
| Corduliidae | 2 | 1 | 1 | WC<sup>8</sup> |
| Culicidae | 2 | - |  |  |
| Lepidoptera | 2 | - |  |  |
| Lepidostomatidae | 2 | 2 |  |  |
| Philopotamidae | 2 | 2 |  |  |
| Synlestidae | 2 | 5 | 3 | EC<sup>8</sup> |
| Trematoda | 2 | - |  |  |
| Aeshnidae | 1 | 4 | 6 | EC<sup>8</sup> |
| Ancyliidae | 1 | 2 |  |  |
| Barbarochthonidae | 1 | 1 |  |  |
| Blephariceridae | 1 | 1 | 2 | EC<sup>15</sup> |
| Bulinidae | 1 | - |  |  |
| Chlorocyphidae | 1 | - | 2 | EC<sup>8</sup> |
| Dixidae | 1 | 1 | 3 | SA<sup>15</sup> |
| Ecnomidae | 1 | 2 |  |  |
| Empididae | 1 | - |  |  |
| Gerridae | 1 | 2 |  |  |
| Glossosomatidae | 1 | 1 |  |  |
| Heptageniidae | 1 | 1 |  |  |
| Hydrophilidae | 1 | 3 |  |  |
| Hydroptilidae | 1 | 4 |  |  |
| Libellulidae | 1 | 8 | 10 | EC<sup>8</sup> |
| Pleidae | 1 | 1 |  |  |
| Syrphidae | 1 | - |  |  |
| Tabanidae | 1 | 1 |  |  |
| Tricorythidae | 1 | 1 |  |  |
| Veliidae | 1 | 1 |  |  |

Citations: <sup>1</sup>Harrison 2004, <sup>2</sup>van Hoven and Day 2002, <sup>3</sup>de Moor 2002, <sup>4</sup>Barber-James and Lugo-Ortiz 2003, <sup>5</sup>Cumberlidge and Daniels 2008, <sup>6</sup>Stevens and Picker 2003, <sup>7</sup>Jansen van Rensburg and Day 2002, <sup>8</sup>Samways and Wilmot 2003, <sup>9</sup>Stals 2008, <sup>10</sup>de Moor and Scott 2003, <sup>11</sup>de Meillon and Wirth 2002, <sup>12</sup>Biström 2008, <sup>13</sup>Nelson 2008, <sup>14</sup>Endrödy-Younga & Stals 2008, <sup>15</sup>Harrison, Prins & Day 2002

When the number of taxa are compared across methods for the seven sites (Fig. 7), the SASS-level taxa for both morphology and DNA-based methods are similar. Species level assignment was poor as the reference library for South African macroinvertebrate fauna is limited, and only 42 out of 254 morphotaxa and 57 out of 404 mOTUs could be assigned to species. If the mOTU levels are considered without assignment, then the spread of mOTU-taxa is more apparent and mean mOTU numbers found with each method are considerably more variable (Fig. 7), highlighting the increased information obtainable at a higher taxon resolution.

**Fig. 7.**
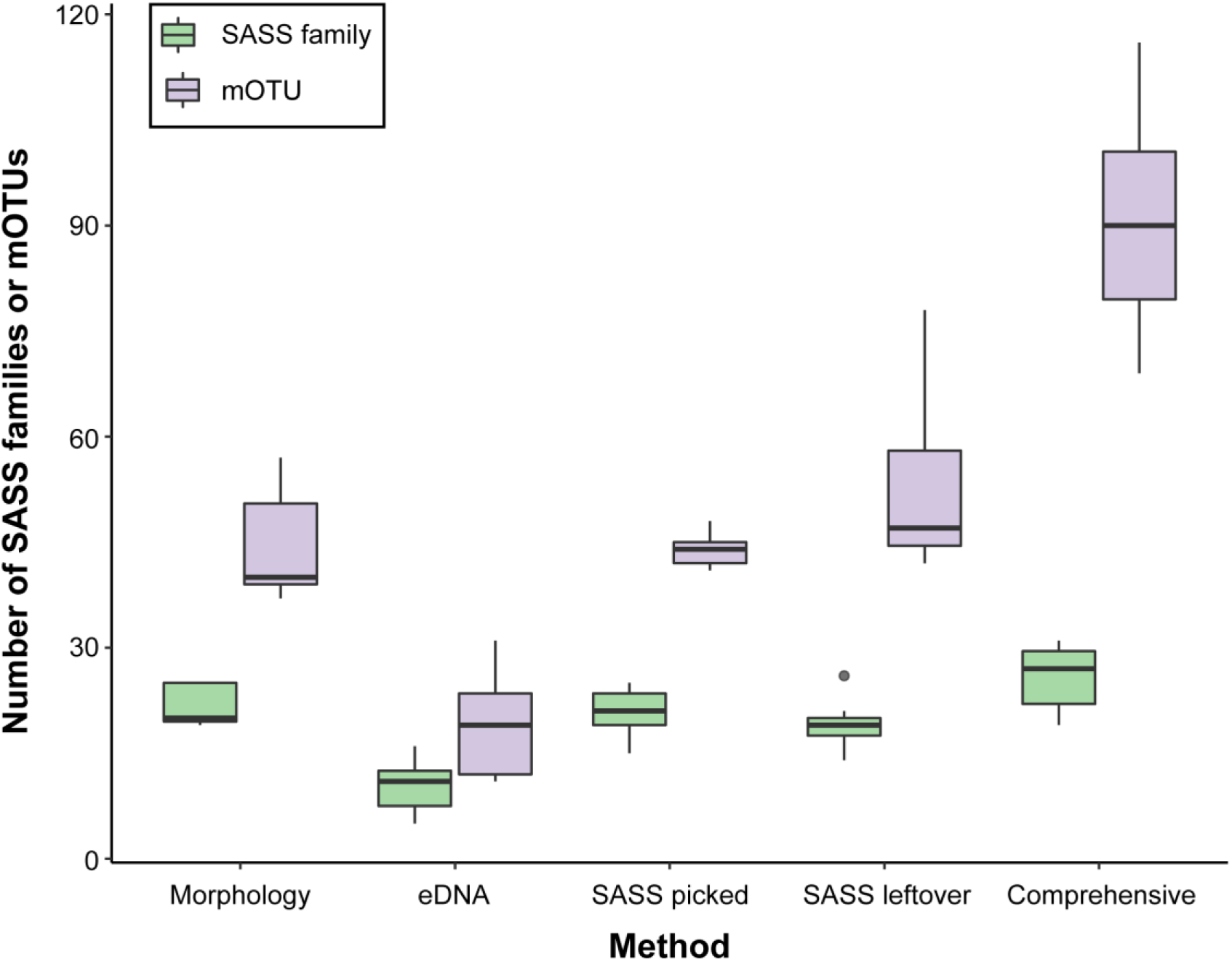
Box and whisker plot showing the mean and interquartile range for SASS-level (green) and OTU-level (purple) taxon numbers for each method over the seven sites.

### mOTU recovery compared to historical records (Table 1, Fig. S3)

Comparing mOTUs in these three rivers against historical records of species from the region highlighted several groups with potentially high levels of cryptic diversity (Table 1). This was especially apparent within the true flies (Diptera) where molecular methods found a total of 101 mOTUs for Chironomidae (Comprehensive = 88; SASS picked = 15; SASS leftover = 65; eDNA = 21) whereas current records from these rivers include two species, with an estimated 30 species from the wider Eastern Cape province. Within the Simuliidae (Diptera) molecular methods found 34 mOTUs (Comprehensive = 31; SASS picked = 23; SASS leftover = 29; eDNA = 11) whereas current records from these rivers include 12 species, with 23 species known from the wider Eastern Cape province. Within the mayflies (Ephemeroptera) two families showed high levels of cryptic diversity; in the Baetidae (Ephemeroptera) molecular methods found 24 mOTUs (Comprehensive = 24; SASS picked = 23; SASS leftover = 19; eDNA = 8) whereas current records from these rivers include 17 species, with 37 species known from the entire region of southern Africa. Whereas within the Leptophlebiidae (Ephemeroptera) molecular methods found 9 mOTUs (Comprehensive = 8; SASS picked = 9; SASS leftover = 8; eDNA = 4) whereas current records from these rivers include five species, with 20 species known from the entire region of southern Africa. Within the stoneflies (Plecoptera) the Notonemouridae exhibited high levels of cryptic diversity with the molecular methods finding 13 mOTUs (Comprehensive = 13; SASS picked = 7; SASS leftover = 7; eDNA = 1) whereas current records from these rivers include 2 species, with 14 species known from South Africa.

Within the dragonflies and damselflies (Odonata) the family Platycnemididae exhibited very high levels of cryptic diversity with the molecular methods finding 10 mOTUs (Comprehensive = 5; SASS picked = 6; SASS leftover = 2; eDNA = 0) whereas current records from these rivers include one species, with only two species known from the wider Eastern Cape province. Within the Oligochaeta which are typically not identified further in the SASS protocol, molecular methods found 42 mOTUs (Comprehensive = 34; SASS picked = 8; SASS leftover = 28; eDNA = 2) whereas current records from these rivers include one species, with 29 species known from the wider Eastern Cape province. Within the Potamonautidae (Crustacea) molecular methods found 23 mOTUs (Comprehensive = 22; SASS picked = 4; SASS leftover = 3; eDNA = 2) whereas current records from these rivers include 2 species, with 14 species known from South Africa.

## Discussion

### DNA methods comparisons

In this study, we compared four DNA-based methods for recovering biodiversity at mixed higher taxon (typically family level) and the mOTU level in the context of water quality biomonitoring and rapid biodiversity assessment. The methods included a standardised SASS sample split into the picked individuals (SASS picked) and the leftover debris (SASS leftover), a Comprehensive sample searching for a longer time period with a finer mesh size, and an eDNA sample of filtered water from each site. Although the SASS picked samples contained mostly target sequences as a result from the sorting and picking process (Fig. 2), this method also recovered a lower diversity at the mOTU level than the SASS leftover debris which was part of the same sample (Fig. 3 and 7). When comparing morphotaxa with the SASS picked mOTUs (essentially the same sample) some of the discrepancies observed may be due to primer bias, rare taxa and the amplification of gut-contents. Other studies have highlighted issues with sorting and picking samples as it is time-consuming and introduces errors associated with missed taxa (Haase et al., 2006, 2010) and is especially difficult to manage for large samples (Bongard, 2011).

The Comprehensive and SASS leftover samples showed similar proportions of non-target reads, and similar compositions of target and non-target taxa detected, however sample inhibition, due to the substantial debris in both of these sample types, was not apparent, although this was not experimentally validated in our study.

The Comprehensive method found the highest number of target taxa and the highest number of unique target taxa, suggesting that a longer sampling time with a smaller size mesh net, followed by whole sample homogenisation captures more of the local macroinvertebrate diversity than either the standardised SASS sampling protocol, or the eDNA sampling method tested in our study. This increase in mOTU diversity recovered in the Comprehensive sample is potentially due to the inclusion of taxa that would morphologically be considered as “out of season” due to their lifestage (e.g. eggs / small larvae) and otherwise undetectable. Thus, the Comprehensive sampling approach has the potential to reduce the number of sampling visits to a site if unseasonal taxa can also be detected, however this needs further testing.

The high diversity recovered from both the SASS leftovers and the Comprehensive samples are promising as the processing time is dramatically reduced if the invertebrates do not need to be removed for identification and more freshwater macroinvertebrate studies are focusing on unsorted whole-sample homogenisation (Andújar et al., 2018; Dowle et al., 2016; Gardham et al., 2014; Hajibabaei et al., 2011; Majaneva et al., 2018).

Our eDNA sampling method found between 2 to 10 times more mOTUs than any other method, however a remarkable proportion of reads were discarded during bioinformatics processing, as many reads were short or non-target reads (e.g. bacteria, Chromista, Plantae and some Chordata). Thus, the number of target taxa detected with eDNA was lower than for all other methods. These non-target reads are likely due to the highly degenerate primers used, which may be unsuitable for eDNA based studies of freshwater macroinvertebrates (Macher et al., 2018; Smith et al., 2012). Although the eDNA samples recovered fewer taxa than the kick-samples, eDNA found more mOTUs for the Gastropoda, Amphipoda, Porifera (sponges), Coelenterata and Platyhelminthes than any other method (Fig. S3) suggesting they may complement traditional kick-sampling methods for these groups.

While these results, supported by a number of other studies, show that eDNA as currently sampled and analysed is an unlikely alternative to sampling whole organisms for detection of whole macroinvertebrate communities in the near term (Barnes & Turner, 2016; Blackman et al., 2019; Hajibabaei et al., 2019; Macher et al., 2018), the field is advancing at a rapid rate (Blackman et al., 2019; Deiner et al., 2017; Deiner, Mächler, et al., 2016; Li et al., 2018; Mächler et al., 2019; Majaneva et al., 2018). Our stringent approach using sequencing replicates should substantially reduce OTUs remaining which are a result of sequencing error but is also likely to have resulted in target taxa being lost from the eDNA samples at this sequencing depth. Nonetheless, in the context of SASS water quality biomonitoring in South Africa our results highlight that eDNA has the potential to be incorporated as it found similar water quality scores to the morphological-based methods in 5 out of 7 sites, however additional research is required to explore macroinvertebrate eDNA metabarcoding across the water quality bands used in South Africa and with more targeted primers. For fish, Pochardt et al (Pochardt, Allen, Hart, Miller, & Yu, 2019) used eDNA to estimate populations over several years, providing an example of advanced developments in eDNA applications to assist with management questions, assessing both presence and abundance of the target organisms. Our results show the potential use of eDNA in assisting with ecosystem management by offering a unique method of biomonitoring, and once established, can provide a much safer and quicker way of monitoring rivers than the current practice which involves physically collecting invertebrate samples from rivers which may harbour dangerous animals such as crocodiles.

### Comparison of methods at family level and water quality metrics

For most biodiverse countries, DNA reference libraries are still a major limiting factor (e.g. Venter & Bezuidenhout, 2016), however most taxa can at least be assigned confidently to family level or below, using current data on BOLD, as was the case for the target taxa in this study suggesting that reference libraries do not hinder the incorporation of DNA metabarcoding into the current SASS protocol.

The results show that while some DNA-based metabarcoding methods detect more diversity than morphology, even at family level, they do not always find the same taxa, as has been reported in other studies (Carew et al., 2013; Clarke, Beard, Swadling, & Deagle, 2017; Elbrecht, Vamos, Meissner, et al., 2017; Gibson et al., 2015; Lejzerowicz et al., 2015; Zimmermann, Glöckner, Jahn, Enke, & Gemeinholzer, 2015). The Comprehensive and SASS-picked samples detected more of the morphologically detected taxa than the SASS leftover samples, which is to be expected as specimens were removed to form the leftover samples. All three kick-net based invertebrate sampling methods detected more target taxa than the eDNA samples. However, despite these differences in taxon recovery between methods at the SASS-level, results for water quality assessment metrics were similar across morphology-based and DNA-based methods. All rivers sampled are considered to be in a good (biological band “B” category) to natural state (biological band “A” category), which was reflected in both the morphology-based and DNA-based results. While these results suggest that metabarcoding (with the exception of eDNA in this case) can produce usable data for current assessment techniques, the DNA methods tested here are able to detect more families than are be detected with morphology, thus their inclusion into the current morphology-based SASS indices could lead to inflated results. While these results are very promising for DNA based water quality assessment in South Africa it is clear that further research at sites which represent the full spectrum of water quality bands are required to facilitate intercalibration of these new DNA methods.

The large variation in numbers and types of taxa found across methods indicates that the family (SASS) level taxon identification is potentially too coarse to make sound assessments on ecological changes in water quality. Our samples differed considerably in terms of taxon scores and numbers, yet when compared at the SASS-level of biological bands, these differences became minimal. A number of studies have warned that family-level identification, being a compromise between accuracy and the level of taxonomic expertise required, is not precise enough to assess water quality changes, primarily due to the different habitat requirements and pollution tolerances found between species within the same family (Barber-James & Pereira-da-Conceicoa, 2016; de Moor, 2002a; Odume, Muller, Arimoro, & Palmer, 2012; Odume, Palmer, Arimoro, & Mensah, 2015). The current lack of data on the habitat requirements and pollution tolerances of some groups can prove problematic for monitoring, especially in biodiverse but water-stressed areas such as South Africa, however a DNA based approach may rapidly provide these data over large taxonomic and geographic scales when combined with relevant information on the sites sampled and the inclusion of reference sites.

Accurate estimation of taxon abundance is one of the current issues facing metabarcoding (Dowle et al., 2016; Elbrecht & Leese, 2015; Elbrecht, Vamos, Meissner, et al., 2017; Piñol, San Andrés, Clare, Mir, & Symondson, 2014), and is hindering practical application for biomonitoring in countries where water quality assessment metrics are required to include abundance measures. Although abundance is recorded and used for other indices in South Africa, it is not used directly for the SASS biomonitoring protocol, and when coupled with the use of family level of identification, would simplify the intercalibration of a DNA metabarcoding method for water quality assessment. The relationship between abundance at the SASS (mixed taxon) level and relative abundance of sequencing reads found a significant positive correlation for combined samples (Fig. 6) with a reasonable adjusted R^2^ value. Other studies have also shown that despite promising significant linear relationships, abundance is not successfully estimated mainly due to poor fit and large differences in magnitude of scatter (e.g. Carew et al., 2013; Clarke et al., 2017; Dowle et al., 2016; Elbrecht, Vamos, Meissner, et al., 2017; Leray & Knowlton, 2017), however additional samples are needed to investigate this relationship and the potential to use sequence reads to estimate abundance in the broad categories currently used by SASS.

Elbrecht et al (2017) provides a detailed overview of factors limiting the application of DNA metabarcoding for ecosystem assessment, and suggestions for potential solutions. Although protocols using DNA metabarcoding for monitoring still need to be further verified and validated, the technique is promising and provides detailed, reproducible and accurate results. Once methods and procedures are standardised, molecular methods are more likely to be integrated into official, routine monitoring programs (Blackman et al., 2019). The metrics used here are designed specifically for the SASS protocol and a mixed-taxon level morphological identification.

### Taxon recovery at higher resolution & mOTU vs historical records

Our results indicate that DNA-based methods offer a higher taxonomic resolution compared to morphology-based identification, in agreement with previous studies (Baird & Sweeney, 2011; Elbrecht, Vamos, Meissner, et al., 2017; Gibson et al., 2015; Stein et al., 2014; Sweeney, Battle, Jackson, & Dapkey, 2011). Taxonomically difficult groups, like Diptera (Chironomidae and Simuliidae in particular), mites and Oligochaeta are often not included at a high taxonomic resolution in biomonitoring protocols, however DNA metabarcoding is a powerful tool in this respect and is able to uncover remarkable mOTU diversity for these groups. Across the sites, 101 chironomid mOTUs were found using the Comprehensive method, while less than a third of that was picked from the SASS samples as morphospecies, and this diversity is usually recorded as a single family and potentially misrepresented by a relatively low taxon score. A similar pattern was seen within nearly all target groups in this study, in many cases a higher number of mOTUs were found than the number of species known from the province or even country, which is surprising given our limited sampling in only three rivers.

While access to morphological expertise has always limited the level to which taxa, and therefore patterns of biodiversity, can be identified, it is argued that the mOTU approach overestimates recognizable species (Clare, Chain, Littlefair, & Cristescu, 2016; Flynn, Brown, Chain, Macisaac, & Cristescu, 2015). This is particularly influenced by retention of rare sequences (singletons) however these are crucial for biodiversity studies (Clare et al., 2016) and despite differences in mOTU recovery rates due to parameter choice, the actual ecological conclusions are not strongly impacted if the error is equal among samples and treatments (Clare et al., 2016). The strength of a mOTU approach is the estimate of diversity ensuring comparability in the absence of described species (Clare et al., 2016) and not originally intended as a species concept (Floyd, Abebe, Papert, & Blaxter, 2002). Nowhere is this more applicable than in biodiversity hotspots, like southern Africa, where species diversity is poorly known. The mOTU approach using DNA-based methods for measuring diversity in areas where species level information is unknown, taxonomic experts and identification keys unavailable, is the most objective method with the added advantage of being more easily standardised. This is especially relevant in cases of cryptic diversity, as community composition data is grossly underestimated using morphological approaches because the largest and most diverse groups (e.g. Chironomidae) are grouped together or omitted from analyses.

The advantage of developing a metabarcoding tool now, despite the limited reference libraries in South Africa, is that the data gathered can be “time stamped” and can be reanalysed as reference libraries are developed in parallel. The mOTU approach can aid in cryptic species detection (Delić, Trontelj, Rendoš, & Fišer, 2017) and rapidly increase the knowledge of species distribution ranges. There is also potential for future assessments using metabarcoding techniques to incorporate and standardise mOTUs (Blaxter et al., 2005) which have not been assigned to the species level, providing a relative diversity with an expected reference condition within an ecoregion. A standardised approach will allow for analysis of the effects of single or multiple stressors for even cryptic and undescribed species (Beermann et al., 2018; Macher et al., 2016). For those genera and species that can be identified using DNA, assessments can be further optimised by including and integrating species functional diversity, ecological preferences and the effects of ecosystem stressors, especially for indicator taxa (Macher et al., 2016).

Although there is still much uncertainty and concern with the conceptual and practical difficulties of translating mOTUs to species (Brown, Chain, Crease, Macisaac, & Cristescu, 2015), the next step of standardising methods to produce measures of biodiversity that are comparable at the mOTU level will be the core of future biodiversity studies, especially where taxonomy is poorly known. It is unrealistic to wait for reference libraries to be built first. DNA metabarcoding approaches to characterise biodiversity have the potential to inspire and focus taxonomic works in hyperdiverse areas, without knowing species level information. Results can be calibrated and improved from existing assessment metrics and reference samples can be kept and assessed for quality control. These mOTU community-level assessments can be associated with ecological trait and preference data based on collection site data, which can be developed into a monitoring database over time. As DNA reference libraries are developed, these datasets will be available for reanalysis and method refinement for better monitoring practice. This detailed level of information can then be used for biomonitoring, biodiversity and ecological impact assessments, providing detailed evidence that can be used to inform managers, governments and policy.

## Conclusions

DNA techniques, especially those using whole sample homogenisation, are not only able to recover the same water quality assessments as morphological SASS, but are able to identify taxa to a much higher resolution. Intensive search comprehensive sampling out-performed all other methods for the number of taxa detected, highlighting that the standardised SASS sampling method does miss relevant target taxa present in the environment, potentially due to the larger mesh size or limited sampling time. Comprehensive sampling with whole-sample homogenisation not only offers the potential to drastically reduce sample processing time, but also potentially collects out-of-season taxa.

While Comprehensive sample homogenisation is a method that has demonstrated promising results, other techniques using fixatives from bulk samples are being developed (Blackman et al., 2019; Carew, Coleman, & Hoffmann, 2018; Erdozain et al., 2019; Martins et al., 2019; Zizka, Leese, Peinert, & Geiger, 2019) and would be advantageous in that the sample remains intact for morphological identification if needed. The strength of eDNA methods for whole catchment assessments is promising (Blackman et al., 2019) and has the potential to benefit biomonitoring protocols by detecting different organisms and offering a safer alternative for assessing dangerous rivers.

This study shows that unsorted samples, including those with significant debris recover the same water quality scores as a morphology-based assessment and recovers much higher diversity scores than both picked and eDNA samples. This approach has significant advantages in that it provides both water quality assessment and species level data at the same time. Thereby making it possible to integrate DNA-based approaches into existing metrics quickly while providing much more information for the development of more refined metrics at the species or mOTU level with distributional data which, as reference libraries are developed, can be used for conservation and biodiversity management.

## Author contributions

LP and BP designed the study. LP, HB-J and BP carried out sample collection. LP, BP, HB-J, AH and AB generated the data. LP, VE and AB analysed the data. LP, VE and BP wrote the original manuscript, all authors contributed to revisions and accepted the final version.

## Data Archiving Statement

Raw MiSeq data can be found here: https://doi.org/10.5281/zenodo.3462633

## Supporting Information

Fig. S1: South African Scoring System (SASS) scoring sheet

Fig. S2. Map showing rivers and sites

Table S1. Morphologically identified taxa at SASS level (presence/absence)

Fig. S3. Detailed barplot showing the number of mOTUs per DNA-based method and morphotaxa found for each SASS family

## Acknowledgements

This research was funded by the Department of Business, Energy & Industrial Strategy Rutherford Fund. Florian Leese, Edith Vamos, Cristina Hartmann and Chris Hempel are thanked for providing the indexed BF2 / BR2 primers. Steve Russell (NHM) is thanked for barcoding the voucher specimens. The National Research Foundation (NRF), South Africa, incentive funding grant number 85286, and Rhodes University Research Council are also acknowledged for funding. Many thanks to Albany Museum staff who helped us in the field, Alex Holland, Musa Mlambo, Nonkazimulo Mdidimba, Ina Ferreira and Zezethu Mnqeta. Rhodes University students, Lauren James and Jacqui James. Juan Tedder (GroundTruth), Kirsty Venter and James Pereira da Conceicoa. Special thanks to Dirk Roux and Nerina Kruger (SANPARKS) for assisting with permits and accommodation and Melanie de Morne (SANPARKS) for assisting in the field. Research material was collected under a once off sampling permit for the Tsitsikamma Section - Garden Route National Park issued on 2017/11/10 by Nerina Kruger, permit number 0056_AAA041-0053 and in the Eastern Cape under CRO 69/17CR and CRO 70/17CR.

