## Supplementary files for "Metabarcoding unsorted kick-samples facilitates macroinvertebrate-based biomonitoring with increased taxonomic resolution, while outperforming environmental DNA"

### SASS Version 5 Score Sheet

**Date:** / /200__

**Collector:** . .

Grid Reference: WGS-84 | Cape datum

S: ° ' , " E: ° ' , "

**Site code:**

**River:...** ... .... ... ........................

Site description:…..........….………...............

Weather Condition:…...….….….…..............

Temp:......…...….°C pH:……….....….....

DO:……….........…mg/l Cond:…...……mS/m

**Biotopes sampled:**

SIC ...................... Time..................minutes

SOOC.................. Time..................minutes

Average size of stones………………….cm

Bedrock...............

Aquatic veg'n......... Dom. sp.................................

MvegIC…………. Dom. sp.................................

MvegOOC………. Dom. sp.................................

Gravel.....…......... Sand.................

Mud.......…...........

Hand picking/Visual observation…………….

**Flow:** Low/Medium/High/Flood

**Turbidity:** Low/Medium/High

**Riparian land use:**

**Disturbance in the river:** eg. sandwinning, cattle drinking point, floods etc.

**Observations:** eg. smell and colour of water, petroleum, dead fish, etc.

| Taxon | | S | Veg | GSM | TOT |
|---|---|---|---|---|---|
| **PORIFERA** | 5 | | | | |
| **COELENTERATA** | 1 | | | | |
| **TURBELLARIA** | 3 | | | | |
| **ANNELIDA** | | | | | |
| Oligochaeta | 1 | | | | |
| Leeches | 3 | | | | |
| **CRUSTACEA** | | | | | |
| Amphipoda | 13 | | | | |
| Potamonautidae* | 3 | | | | |
| Atyidae | 8 | | | | |
| Palaemonidae | 10 | | | | |
| **HYDRACARINA** | 8 | | | | |
| **PLECOPTERA** | | | | | |
| Notonemouridae | 14 | | | | |
| Perlidae | 12 | | | | |
| **EPHEMEROPTERA** | | | | | |
| Baetidae 1sp | 4 | | | | |
| Baetidae 2 sp | 6 | | | | |
| Baetidae > 2 sp | 12 | | | | |
| Caenidae | 6 | | | | |
| Ephemeridae | 15 | | | | |
| Heptageniidae | 13 | | | | |
| Leptophlebiidae | 9 | | | | |
| Oligoneuridae | 15 | | | | |
| Polymitarcyidae | 10 | | | | |
| Prosopistomatidae | 15 | | | | |
| Teloganodidae SWC | 12 | | | | |
| Tricorythidae | 9 | | | | |
| **ODONATA** | | | | | |
| Calopterygidae ST,T | 10 | | | | |
| Chlorocyphidae | 10 | | | | |
| Chlorolestidae | 8 | | | | |
| Coenagrionidae | 4 | | | | |
| Lestidae | 8 | | | | |
| Platycnemidae | 10 | | | | |
| Protoneuridae | 8 | | | | |
| Aeshnidae | 8 | | | | |
| Corduliidae | 8 | | | | |
| Gomphidae | 6 | | | | |
| Libellulidae | 4 | | | | |
| **LEPIDOPTERA** | | | | | |
| Pyralidae | 12 | | | | |

| Taxon | | S | Veg | GSM | TOT |
|---|---|---|---|---|---|
| **HEMIPTERA** | | | | | |
| Belostomatidae* | 3 | | | | |
| Corixidae* | 3 | | | | |
| Gerridae* | 5 | | | | |
| Hydrometridae* | 6 | | | | |
| Naucoridae* | 7 | | | | |
| Nepidae* | 3 | | | | |
| Notonectidae* | 3 | | | | |
| Pleidae* | 4 | | | | |
| Veliidae/M...veliidae* | 5 | | | | |
| **MEGALOPTERA** | | | | | |
| Corydalidae | 8 | | | | |
| Sialidae | 6 | | | | |
| **TRICHOPTERA** | | | | | |
| Dipseudopsidae | 10 | | | | |
| Ecnomidae | 8 | | | | |
| Hydropsychidae 1 sp | 4 | | | | |
| Hydropsychidae 2 sp | 6 | | | | |
| Hydropsychidae > 2 sp | 12 | | | | |
| Philopotamidae | 10 | | | | |
| Polycentropodidae | 12 | | | | |
| Psychomyiidae/Xiphocen | 8 | | | | |
| **Cased caddis:** | | | | | |
| Barbarochthonidae SWC | 13 | | | | |
| Calamoceratidae ST | 11 | | | | |
| Glossosomatidae SWC | 11 | | | | |
| Hydroptilidae | 6 | | | | |
| Hydrosalpingidae SWC | 15 | | | | |
| Lepidostomatidae | 10 | | | | |
| Leptoceridae | 6 | | | | |
| Petrothrincidae SWC | 11 | | | | |
| Pisuliidae | 10 | | | | |
| Sericostomatidae SWC | 13 | | | | |
| **COLEOPTERA** | | | | | |
| Dytiscidae* | 5 | | | | |
| Elmidae/Dryopidae* | 8 | | | | |
| Gyrinidae* | 5 | | | | |
| Haliplidae* | 5 | | | | |
| Helodidae | 12 | | | | |
| Hydraenidae* | 8 | | | | |
| Hydrophilidae* | 5 | | | | |
| Limnichidae | 10 | | | | |
| Psephenidae | 10 | | | | |

| Taxon | | S | Veg | GSM | TOT |
|---|---|---|---|---|---|
| **DIPTERA** | | | | | |
| Athericidae | 10 | | | | |
| Blepharoceridae | 15 | | | | |
| Ceratopogonidae | 5 | | | | |
| Chironomidae | 2 | | | | |
| Culicidae* | 1 | | | | |
| Dixidae* | 10 | | | | |
| Empididae | 6 | | | | |
| Ephydridae | 3 | | | | |
| Muscidae | 1 | | | | |
| Psychodidae | 1 | | | | |
| Simuliidae | 5 | | | | |
| Syrphidae* | 1 | | | | |
| Tabanidae | 5 | | | | |
| Tipulidae | 5 | | | | |
| **GASTROPODA** | | | | | |
| Ancylidae | 6 | | | | |
| Bulininae* | 3 | | | | |
| Hydrobiidae* | 3 | | | | |
| Lymnaeidae* | 3 | | | | |
| Physidae* | 3 | | | | |
| Planorbinae* | 3 | | | | |
| Thiaridae* | 3 | | | | |
| Viviparidae* ST | 5 | | | | |
| **PELECYPODA** | | | | | |
| Corbiculidae | 5 | | | | |
| Sphaeriidae | 3 | | | | |
| Unionidae | 6 | | | | |
| **SASS Score** | | | | | |
| **No. of Taxa** | | | | | |
| **ASPT** | | | | | |

Sample collection effort exceeds method? .....

**Other biota including juveniles:**

**Comments:**

Procedure: 'Kick SIC & bedrock for 2 mins, max. 5 mins; Kick SOOC & bedrock for 1 min; Sweep marginal vegetation (IC & OOC) for 2m total and aquatic veg 1m$^2$; Stir & sweep gravel, sand, mud for 1 min total; * = airbreathers; Hand picking & visual observation for 1 min — record in biotope where found; Score for 15 mins/biotope but stop if no new taxa seen after 5 mins; 'Estimate abundances: 1 = 1, A = 2–10, B = 10–100, C = 100–1 000, D = >1 000; S = Stone, rock & solid objects; Veg = All vegetation; GSM = Gravel, sand, mud; SWC = South Western Cape; T = Tropical; ST = Sub-tropical; Rate each biotope sampled: 1 = very poor (i.e. limited diversity), 5 = highly suitable (i.e. wide diversity)

Fig. S1: South African Scoring System (SASS) scoring sheet

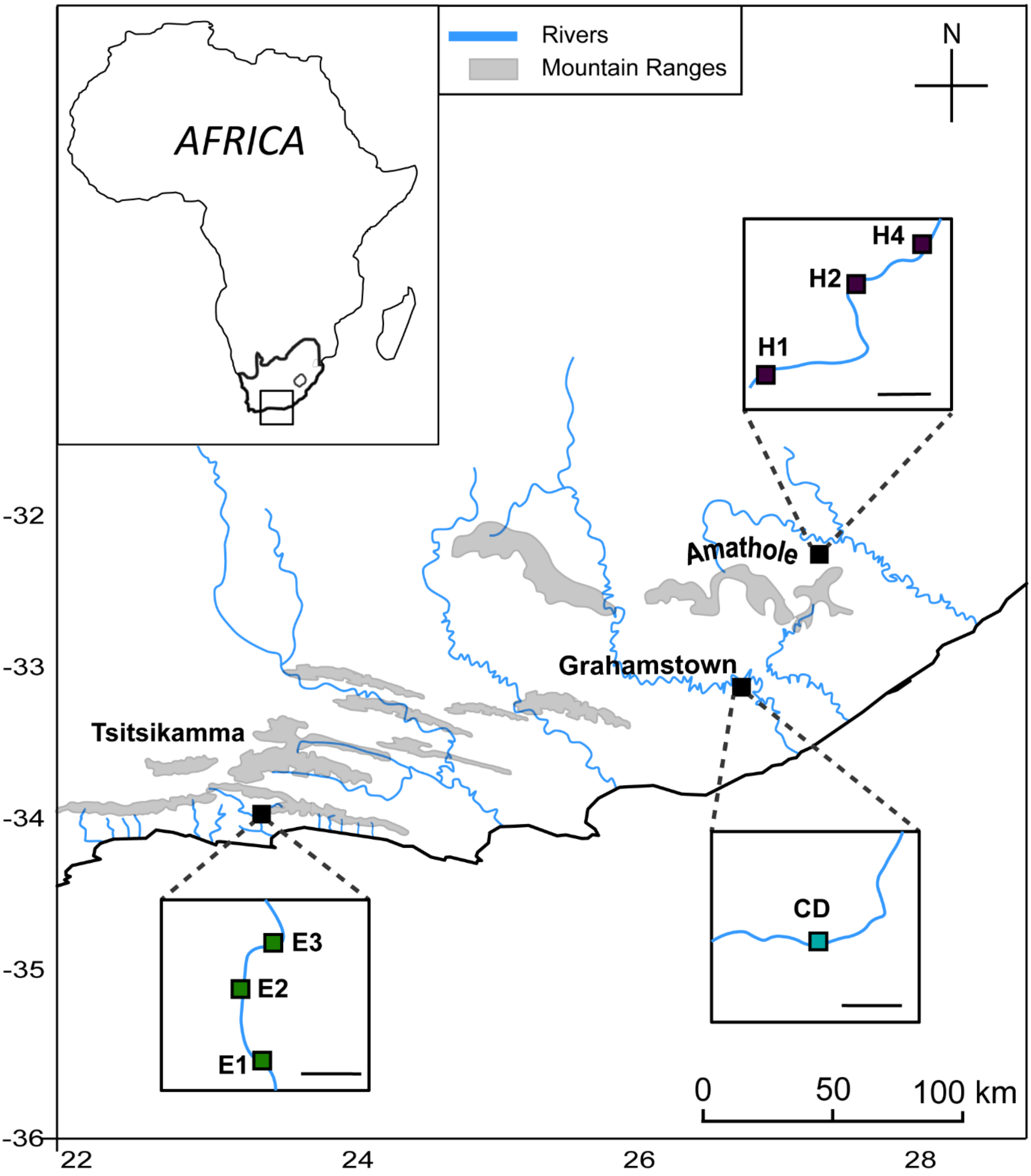


Figure S2. Sample sites the Elandsbos River in the Tsitsikamma region, a tributary of the Tyume River (Hobbiton's "Eerste" stream) in Hogsback of the Amathole mountain region and the Berg River at Craigdoone near Grahamstown. Each inset map of sites has a scale bar = 100m

Table S3. Morphologically identified taxa at SASS level (presence/absence)

| M_ID | phylum | class/subclass | order | family | taxa | E1 | E2 | E3 | CD | H1 | H2 | H4 | Taxonomic rank |
|---|---|---|---|---|---|---|---|---|---|---|---|---|---|
| 1 | Platyhelminthes | | | | Platyhelminthes | 0 | 0 | 0 | 0 | 0 | 0 | 1 | Phylum |
| 2 | Annelida | Clitellata/Oligochaeta | | | Oligochaeta | 0 | 1 | 1 | 1 | 0 | 0 | 1 | subclass |
| 3 | Annelida | Clitellata/Hirudinea | | | Hirudinea | 0 | 0 | 0 | 0 | 1 | 0 | 0 | subclass |
| 4 | Arthropoda | Malacostraca | Amphipoda | | Amphipoda | 0 | 0 | 1 | 0 | 0 | 0 | 0 | order |
| 5 | Arthropoda | Malacostraca | Decapoda | Potamonautidae | Potamonautidae | 0 | 0 | 1 | 1 | 1 | 0 | 0 | family |
| 6 | Arthropoda | Insecta | Plecoptera | Notonemouridae | Notonemouridae | 1 | 1 | 1 | 0 | 1 | 1 | 1 | family |
| 7 | Arthropoda | Insecta | Ephemeroptera | Baetidae | Baetidae | 1 | 1 | 1 | 1 | 1 | 1 | 1 | family |
| 8 | Arthropoda | Insecta | Ephemeroptera | Caenidae | Caenidae | 1 | 0 | 0 | 1 | 1 | 0 | 0 | family |
| 9 | Arthropoda | Insecta | Ephemeroptera | Leptophlebiidae | Leptophlebiidae | 1 | 1 | 1 | 1 | 1 | 1 | 1 | family |
| 10 | Arthropoda | Insecta | Ephemeroptera | Teloganodidae | Teloganodidae | 1 | 1 | 1 | 0 | 1 | 1 | 1 | family |
| 11 | Arthropoda | Insecta | Ephemeroptera | Tricorythidae | Tricorythidae | 0 | 0 | 0 | 0 | 1 | 1 | 1 | family |
| 12 | Arthropoda | Insecta | Odonata | Synlestidae | Synlestidae | 0 | 0 | 1 | 1 | 1 | 0 | 0 | family |
| 13 | Arthropoda | Insecta | Odonata | Coenagrionidae | Coenagrionidae | 1 | 1 | 1 | 1 | 0 | 0 | 0 | family |
| 14 | Arthropoda | Insecta | Odonata | Platycnemididae | Platycnemididae | 0 | 1 | 1 | 0 | 0 | 0 | 0 | family |
| 15 | Arthropoda | Insecta | Odonata | Aeshnidae | Aeshnidae | 0 | 0 | 1 | 0 | 0 | 0 | 0 | family |
| 16 | Arthropoda | Insecta | Odonata | Gomphidae | Gomphidae | 0 | 0 | 0 | 1 | 0 | 0 | 0 | family |
| 17 | Arthropoda | Insecta | Odonata | Libellulidae | Libellulidae | 0 | 0 | 0 | 0 | 0 | 0 | 0 | family |
| 18 | Arthropoda | Insecta | Lepidoptera | Pyralidae | Pyralidae | 0 | 0 | 0 | 0 | 0 | 0 | 1 | family |
| 19 | Arthropoda | Insecta | Hemiptera | Corixidae | Corixidae | 1 | 0 | 0 | 0 | 0 | 0 | 0 | family |
| 20 | Arthropoda | Insecta | Hemiptera | Gerridae | Gerridae | 0 | 0 | 0 | 0 | 0 | 0 | 0 | family |
| 21 | Arthropoda | Insecta | Hemiptera | Naucoridae | Naucoridae | 1 | 1 | 1 | 0 | 0 | 0 | 0 | family |
| 22 | Arthropoda | Insecta | Hemiptera | Notonectidae | Notonectidae | 1 | 0 | 1 | 0 | 0 | 0 | 0 | family |
| 23 | Arthropoda | Insecta | Hemiptera | Veliidae | Veliidae | 1 | 1 | 0 | 0 | 0 | 1 | 0 | family |
| 24 | Arthropoda | Insecta | Megaloptera | Corydalidae | Corydalidae | 1 | 1 | 1 | 0 | 1 | 1 | 1 | family |
| 25 | Arthropoda | Insecta | Trichoptera | Ecnomidae | Ecnomidae | 1 | 0 | 1 | 0 | 0 | 0 | 0 | family |
| 26 | Arthropoda | Insecta | Trichoptera | Hydropsychidae | Hydropsychidae | 0 | 0 | 0 | 1 | 1 | 1 | 1 | family |
| 27 | Arthropoda | Insecta | Trichoptera | Philopotamidae | Philopotamidae | 0 | 1 | 1 | 1 | 0 | 0 | 0 | family |
| 28 | Arthropoda | Insecta | Trichoptera | Psychomyiidae | Psychomyiidae | 0 | 1 | 0 | 0 | 0 | 0 | 0 | family |
| 29 | Arthropoda | Insecta | Trichoptera | Barbarochthonidae | Barbarochthonidae | 1 | 1 | 1 | 0 | 0 | 0 | 0 | family |
| 30 | Arthropoda | Insecta | Trichoptera | Glossosomatidae | Glossosomatidae | 1 | 1 | 1 | 0 | 0 | 0 | 0 | family |
| 31 | Arthropoda | Insecta | Trichoptera | Hydroptilidae | Hydroptilidae | 0 | 0 | 0 | 1 | 0 | 0 | 1 | family |
| 32 | Arthropoda | Insecta | Trichoptera | Lepidostomatidae | Lepidostomatidae | 0 | 0 | 0 | 0 | 1 | 1 | 0 | family |

| | | | | | | | | | | | | | |
|---|---|---|---|---|---|---|---|---|---|---|---|---|---|
| 33 | Arthropoda | Insecta | Trichoptera | Leptoceridae | Leptoceridae | 1 | 1 | 1 | 0 | 1 | 1 | 1 | family |
| 34 | Arthropoda | Insecta | Trichoptera | Pisuliidae | Pisuliidae | 1 | 0 | 1 | 0 | 1 | 1 | 0 | family |
| 35 | Arthropoda | Insecta | Trichoptera | Sericostomatidae | Sericostomatidae | 1 | 0 | 0 | 0 | 1 | 1 | 1 | family |
| 36 | Arthropoda | Insecta | Coleoptera | Dytiscidae | Dytiscidae | 0 | 0 | 1 | 1 | 0 | 0 | 0 | family |
| 37 | Arthropoda | Insecta | Coleoptera | Elmidae | Elmidae | 1 | 1 | 0 | 0 | 1 | 0 | 0 | family |
| 38 | Arthropoda | Insecta | Coleoptera | Gyrinidae | Gyrinidae | 1 | 0 | 1 | 1 | 0 | 0 | 0 | family |
| 39 | Arthropoda | Insecta | Coleoptera | Scirtidae | Scirtidae | 1 | 1 | 1 | 0 | 0 | 0 | 1 | family |
| 40 | Arthropoda | Insecta | Diptera | Athericidae | Athericidae | 1 | 0 | 1 | 0 | 0 | 0 | 0 | family |
| 41 | Arthropoda | Insecta | Diptera | Blephariceridae | Blephariceridae | 0 | 0 | 0 | 0 | 1 | 1 | 1 | family |
| 42 | Arthropoda | Insecta | Diptera | Ceratopogonidae | Ceratopogonidae | 0 | 0 | 0 | 0 | 0 | 0 | 1 | family |
| 43 | Arthropoda | Insecta | Diptera | Chironomidae | Chironomidae | 1 | 1 | 1 | 1 | 1 | 1 | 1 | family |
| 44 | Arthropoda | Insecta | Diptera | Culicidae | Culicidae | 1 | 0 | 0 | 1 | 0 | 0 | 0 | family |
| 45 | Arthropoda | Insecta | Diptera | Dixidae | Dixidae | 0 | 0 | 0 | 0 | 1 | 0 | 0 | family |
| 46 | Arthropoda | Insecta | Diptera | Simuliidae | Simuliidae | 1 | 1 | 0 | 1 | 1 | 1 | 1 | family |
| 47 | Arthropoda | Insecta | Diptera | Tabanidae | Tabanidae | 0 | 0 | 0 | 1 | 0 | 0 | 0 | family |
| 48 | Arthropoda | Insecta | Diptera | Tipulidae | Tipulidae | 0 | 0 | 0 | 1 | 1 | 1 | 0 | family |
| 49 | Mollusca | Gastropoda | Hygrophila | Planorbidae | Planorbidae | 0 | 0 | 0 | 1 | 1 | 1 | 1 | family |
| 50 | Mollusca | Gastropoda | Hygrophila | Planorbinae | Planorbinae | 0 | 0 | 0 | 0 | 1 | 0 | 0 | family |
| 51 | Mollusca | Bivalvia | Veneroida | Sphaeriidae | Sphaeriidae | 0 | 0 | 0 | 0 | 1 | 1 | 0 | family |

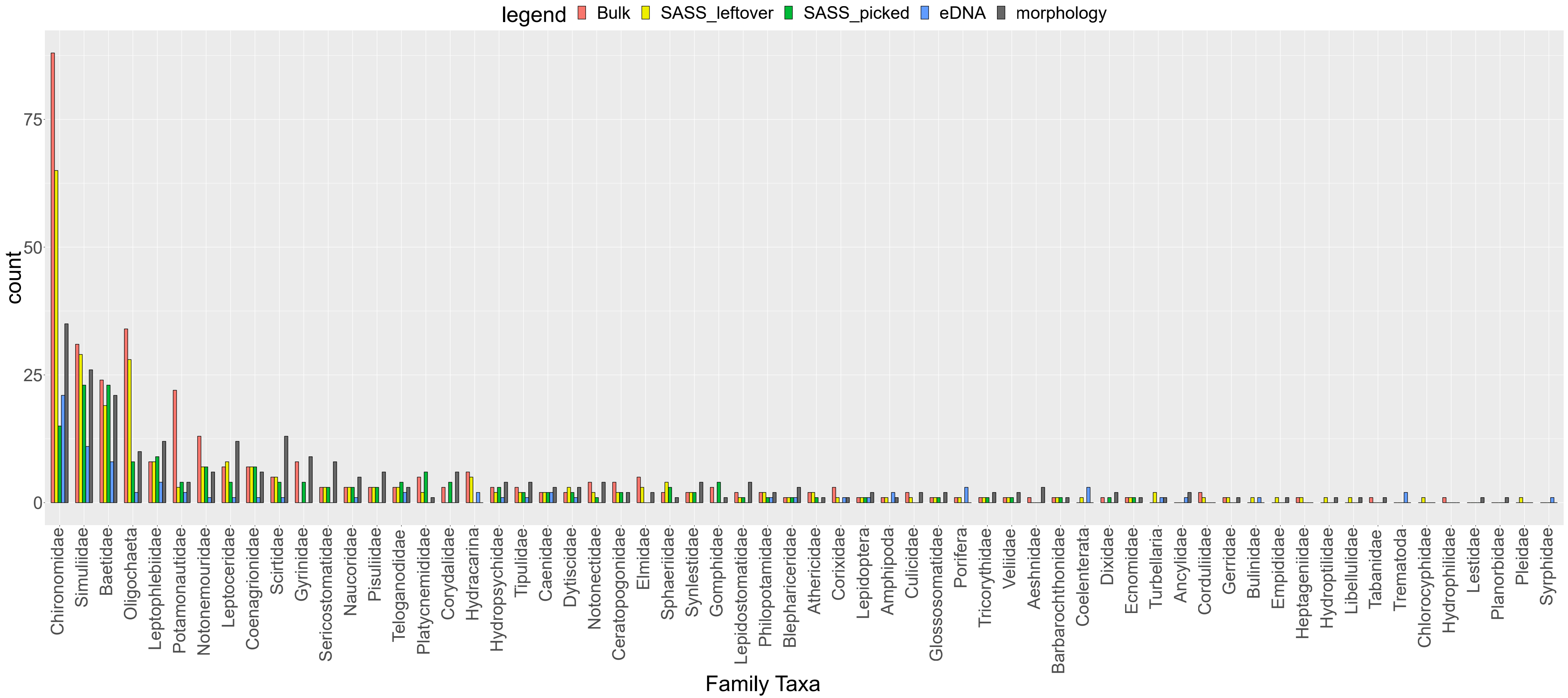


Fig. S3. Detailed barplot showing the number of mOTUs per DNA-based method and morphotaxa found for each SASS family
